# A Genome-Wide CRISPR Screen Identifies Sortilin as the Receptor Responsible for Galectin-1 Lysosomal Trafficking

**DOI:** 10.1101/2024.01.03.574113

**Authors:** Justin Donnelly, Roarke A. Kamber, Simon Wisnovsky, David S. Roberts, Egan L. Peltan, Michael C. Bassik, Carolyn R. Bertozzi

**Affiliations:** Sarafan ChEM-H, Stanford University, Stanford, CA 94305; Department of Chemistry, Stanford University, Stanford, CA 94305; Department of Genetics, Stanford University, Stanford, CA 94305; Department of Anatomy and Bakar Aging Research Institute, University of California at San Francisco, San Francisco, CA 94143; Faculty of Pharmaceutical Sciences, University of British Columbia, Vancouver, Canada, V6T 1Z3; Department of Chemical and Systems Biology, Stanford University, Stanford, CA 94305

## Abstract

Galectins are a family of mammalian glycan-binding proteins that have been implicated as regulators of myriad cellular processes including cell migration, apoptosis, and immune modulation. Several members of this family, such as galectin-1, exhibit both cell-surface and intracellular functions. Interestingly, galectin-1 can be found in the endomembrane system, nucleus, or cytosol, as well as on the cell surface. The mechanisms by which galectin-1 traffics between cellular compartments, including its unconventional secretion and internalization processes, are poorly understood. Here, we determined the pathways by which exogenous galectin-1 enters cells and explored its capacity as a delivery vehicle for protein and siRNA therapeutics. We used a galectin-1-toxin conjugate, modelled on antibody-drug conjugates, as a selection tool in a genome-wide CRISPR screen. We discovered that galectin-1 interacts with the endosome-lysosome trafficking receptor sortilin in a glycan-dependent manner, which regulates galectin-1 trafficking to the lysosome. Further, we show that this pathway can be exploited for delivery of a functional siRNA. This study sheds light on the mechanisms by which galectin-1 is internalized by cells and suggests a new strategy for intracellular drug delivery via galectin-1 conjugation.

---

Galectin-1 (Gal1) is a member of the galectin family of proteins that binds β-galactosides. These are mainly unsialylated *N*-acetyllactosamine (LacNAc) motifs, which are primarily found on N-glycans on cell surfaces or in the extracellular matrix^1–3^. Gal1’s binding partners include integrins^4–6^, CD4^7,8^, CD43^7–10^, CD45^7–9^, and mucins^3,11,12^. The binding of Gal1 to glycans is involved in myriad biological processes, including cell adhesion^4,13^, migration^4,14^, proliferation^14,15^, and apoptosis^10^. Gal1 plays a critical role in the immune system: Gal1 binding to dendritic cells can promote dendritic cell migration and CD4+ T cell proliferation^14^, while its binding to CD43 and CD45 can modulate T cell apoptosis and promote immune tolerance^8,10,16^. Even as one of the most well-studied members of the galectin family, however, many of the specific molecular interactions that give rise to the functional consequences of Gal1 activity have not been fully defined.

Although initially thought to be exclusively localized to the cytoplasm and extracellular matrix, Gal1 has also been shown to translocate into the nucleus under certain conditions^17,18^. In the nucleus, Gal1 has been implicated in regulating gene expression^19^ and RNA splicing^20,21^. Gal1 has been shown to interact with several other nuclear proteins, including pre-mRNA splicing factors like Gemin4^22^, as well as an array of mRNAs^23^, suggesting that it may have as varied roles in nuclear processes as at the cell surface. Moreover, these activities have been shown to have functional importance, directly regulating cell migration^17^, proliferation^19^, and angiogenesis^23,24^.

Evidence exists to support the idea that Gal1 exhibits bidirectional trafficking between the intra- and extracellular spaces through unconventional mechanisms. While it lacks a canonical signal peptide and is synthesized on free cytosolic ribosomes^25^, researchers have shown that Gal1 can be secreted through a Golgi-independent pathway mediated by secretory autophagosomes^26^. Comparatively little is known about the mechanisms by which Gal1 is internalized from the cell surface, though Gal1 synthesized into the ER lumen via docked ribosomes can still reach the cell interior^17^. A better understanding of the specific mechanisms of Gal1 intracellular trafficking could therefore identify methods to control Gal1 localization for research or therapeutic purposes. It could also highlight potential strategies to control the trafficking of endogenous or therapeutic biomolecules, which are of significant interest to drug developers.

Genome-wide CRISPR screening is a powerful tool that enables researchers to identify genes that regulate a particular cellular process or phenotype in an unbiased manner^27,28^. We reasoned that CRISPR screening would be ideally suited to studying the process of Gal1 internalization, as the internalization of Gal1 itself can be used to confer a selectable property onto the cells that take it up. Moreover, the unbiased nature of the screen requires minimal prior knowledge of the mechanistic biology of the process of interest. Here we present a genome-wide screen to identify genes whose knockout ablates Gal1 internalization in live cells. Our study revealed that sortilin is the receptor responsible for sorting Gal1 into the lysosome rather than the rest of the endomembrane system. We leveraged these findings to develop a functional Gal1-siRNA conjugate capable of effecting gene knockdown in a sortilin-dependent manner. Our findings offer new insights into Gal1’s cellular trafficking and suggest potential applications to the delivery of therapeutic siRNAs.

## Results

### Recombinant Gal1 is Internalized in Multiple Cell Lines

In pooled CRISPR screening, a cell line expressing Cas9 is infected with a pooled, genome-wide library of single-guide RNAs (sgRNAs). Cells are then exposed to some type of selective pressure, and sgRNAs that modulate sensitivity to selection become either enriched or depleted compared to initial conditions. Post-hoc analysis of changes in sgRNA abundance through deep sequencing can then be used to identify genes whose disruption modulates the selectable phenotype^27,28^. In order to conduct an effective pooled CRISPR screen, it is thus critical to first design an appropriate selection strategy based on internalization of the Gal1 protein. To this end, we demonstrated that recombinant Gal1 purified from a bacterial source could recapitulate the previously observed internalization of endogenous Gal1. We prepared recombinant Gal1 via heterologous expression in bacteria and confirmed its identity and purity via SDS-PAGE and top-down protein mass spectrometry (TDP-MS). We also confirmed that recombinant Gal1 retained its glycan-binding activity via lactose affinity fast protein liquid chromatography (FPLC) (Supplemental Figure S1).

To assess internalization of recombinant Gal1, we conjugated Gal1 to AlexaFluor-647 (AF647, Supplemental Figure S2), treated cells, and imaged them via confocal microscopy. We observed that Gal1 was internalized into multiple cell lines, including K562, MRC5, and HEK 293T cells. In these cell lines, Gal1 was observed in puncta of varying size and distribution (Figure 1a-c). Next, we used flow cytometry to quantify Gal1 internalization across a broad range of concentrations, and we observed dose-dependent internalization that did not saturate at biologically-relevant concentrations (Figure 1d). To ensure that Gal1 bound to the cell surface did not contribute to these readings, cells were washed several times in PBS with 100 mM lactose before analysis. Importantly, we also used the SYTOX dead cell stain to test whether recombinant Gal1 treatment exhibited intrinsic toxicity, as this could result in selection unrelated to the process of Gal1 internalization. We demonstrated that recombinant Gal1 treatment is non-toxic to K562 cells at concentrations up to 10 µM (Figure 1e-f), which was significantly higher than those necessary to observe significant internalization. These results demonstrate that recombinant Gal1 can suitably recapitulate the internalization of Gal1, enabling the design of a genome-wide CRISPR screen to study this process.

**Figure 1:**
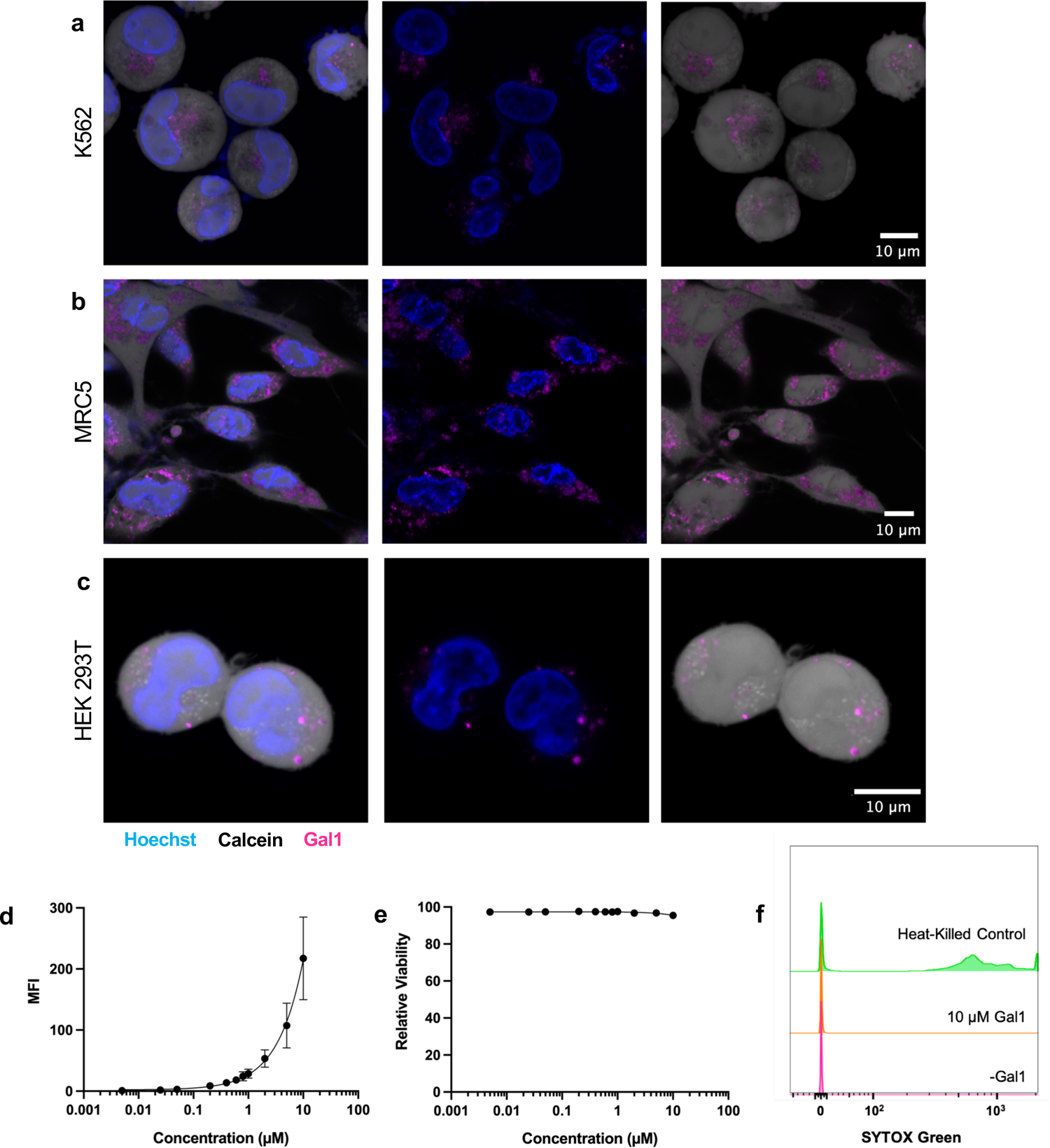
Gal1-AF647 is internalized by multiple cell lines and is non-toxic. **(A)** Confocal images of live K562s ted with Gal1-AF647 (2 µM, magenta) and stained with Hoechst (blue) and Calcein AM (white). **(B)** Confocal ges of live MRC5s treated with Gal1-AF647 (1 µM, magenta) and stained with Hoechst (blue) and Calcein AM te). **(C)** Confocal images of live HEK 293Ts treated with Gal1-AF647 (1 µM, magenta) and stained with chst (blue) and Calcein AM (white). **(D)** Flow cytometry analysis of live K562s treated with a concentration es of Gal1-AF647, then washed with 100 mM lactose before analysis. MFI = mean fluorescence intensity. ts represent averages of two technical and two biological replicates. **(E)** Gal1 toxicity assay. K562s treated with oncentration series of Gal1-AF647 were stained with SYTOX green to measure toxicity. Points represent rages of two technical replicates (error bars not visible). **(F)** Heat-killed cells were used as a positive control to date SYTOX staining of dead cells. Data shown are a representative replicate.

### A Lectin-Drug Conjugate Enables a Gal1 Internalization CRISPR Screen

In order to design a screen to study Gal1 internalization, we elected to use a selection approach based on preferential toxicity. We hypothesized that treatment of a CRISPR screening library with a toxic form of Gal1 would positively select for cells with guides targeting genes necessary for Gal1 internalization (Figure 2a). An analogous approach was recently used to investigate the mechanisms of internalization and toxicity of antibody-drug conjugates (ADCs), which successfully utilized the toxicity of the ADCs themselves as a selection tool. This screen discovered a strong relationship between the toxicity of ADCs and their internalization^29^. To adapt this approach to study the internalization of Gal1, we used Gal1 to develop a biomolecular tool that we term a “lectin-drug conjugate”, or LDC.

**Figure 2:**
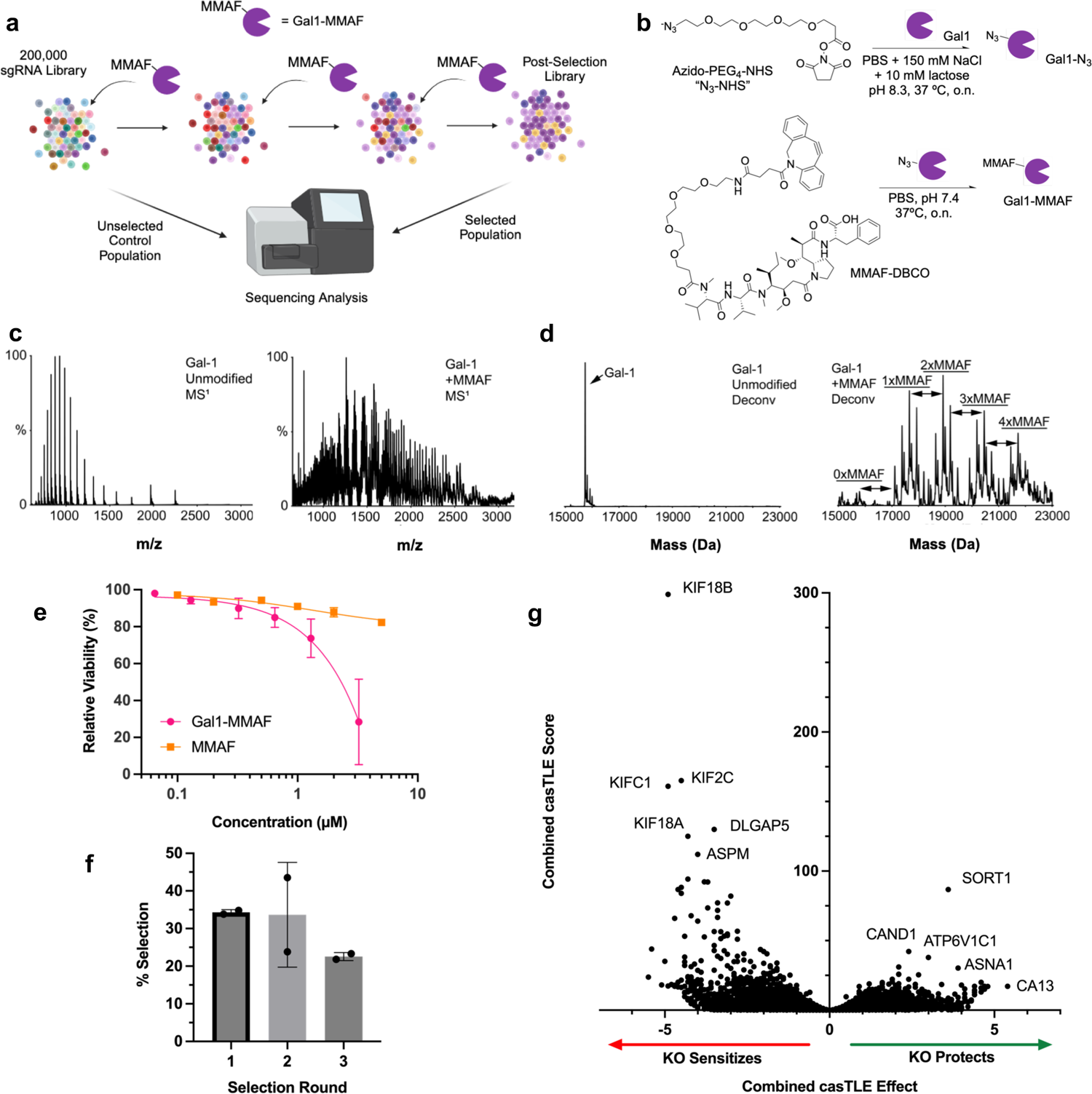
A Genome-Wide CRISPR Screen Identifies Sortilin as a Critical Mediator of Gal1 Internalization in 2 Cells. **(A)** Screen schematic. Three sequential treatments with the LDC Gal1-MMAF enriches Gal1 rnalization-deficient knockouts from a genome-wide knockout library. The relative frequency of gRNA-encoding structs in treated vs. untreated subpopulations is determined via deep sequencing. **(B)** Schematic of LDC aration via functionalization of Gal1 with MMAF. A clickable azide (N_3_) handle is installed on Gal1 using NHS r chemistry (*top*), then functionalized with MMAF-DBCO to yield Gal1-MMAF (*bottom*). The carboxylate group on AF is responsible for its reduced cell permeability relative to MMAE, which lacks this functional group. **(C-D)** Raw and deconvoluted (D) top-down mass spectra of unmodified Gal1 (*left*) and the LDC Gal1-MMAF (*right*). culated DLR = 2.36. **(E)** Flow cytometry analysis of K562 cells treated with Gal1-MMAF or MMAF and then stained SYTOX to measure cell viability. **(F)** Gal1-MMAF treatment-induced selection measured by counting trypan blue-ned cells after each treatment during the CRISPR screen. **(G)** Volcano plot of CRISPR screen sequencing results esenting relative enrichment of knockouts in the Gal1-MMAF-treated population vs control. Genes on the right are more resistant to Gal1-MMAF toxicity than the population as a whole, while genes on the left are more sensitive.

To create our prototypical LDC, we conjugated Gal1 to the ADC toxin MMAF, a highly toxic inhibitor of tubulin polymerization. MMAF is similar to its better-known analogue MMAE, which is often used in commercially-available ADCs, but it has reduced cell permeability thanks to an additional negatively-charged carboxylate group^30^. We prepared the Gal1-MMAF conjugate via copper-free click chemistry^31^, first functionalizing Gal1 with an azide handle and then clicking on MMAF-DBCO (Figure 2b). We then used TDP-MS to characterize Gal1-MMAF and observed a mixture of MMAF-functionalized Gal1 with a distribution of 0 to 4 covalently-attached drug molecules (Figure 2c-d). We determined the overall drug-lectin ratio (DLR) to be 2.36. Next, we tested the LDC’s toxicity in K562 cells. Gal1-MMAF exhibited dose-dependent toxicity and was significantly more toxic than MMAF alone, indicating that its toxicity was dependent on Gal1 internalization (Figure 2e).

We then proceeded to use Gal1-MMAF cytotoxicity to screen against K562 cells transformed with a 200,000-guide CRISPRi gRNA library. We performed three successive treatments with Gal1-MMAF at a concentration close to the LD_50_, each followed by a recovery period. In the third selection round we observed an increase in resistance to Gal1-MMAF selection, indicating the enrichment of a resistant population in response to successive treatments (Figure 2f). Guides from the selected population as well as an untreated control population were then amplified and sequenced. Comparison of the abundance of sgRNA sequences in the treated and untreated populations using the CasTLE algorithm^27^ revealed genes as candidate negative and positive regulators of Gal1-MMAF toxicity. The *SORT1* gene, which encodes the trafficking protein sortilin, was identified as the top positively enriched hit (Figure 2g). A gene ontology (GO) term enrichment of the top 100 positively-enriched hits revealed acidification of endomembrane lumina as a crucial process for Gal1 internalization, with many proton-pumping ATPases and the bicarbonate transporter *SLC4A7* represented. Interestingly, two members of the SCF E3 ligase complex, *CAND1* and *CUL1*, were also positively enriched. Conversely, GO term enrichment of the top 100 negatively-enriched hits revealed processes involved in the toxic mechanism of MMAF, mainly core microtubule functions. Many negatively-enriched hits were also kinesin family (KIF) members, motor proteins that walk along microtubules (Supplemental Figure S3). However, as our goal was to identify cellular machinery involved in promoting Gal1 internalization, and because sortilin is a known trafficking receptor, we focused our follow-up investigations on this protein and its role in Gal1 internalization and trafficking.

### Validation of Sortilin as a Positively-Enriched Hit

Sortilin has been previously shown to mediate internalization of other proteins^32,33^ and serves as a receptor for the signaling peptide neurotensin^34^, but a molecular interaction with Gal1 or a role in its internalization have not yet been reported. In order to validate *SORT1* as a key regulator of Gal1 internalization, we generated a polyclonal sortilin knockout (KO) cell line (Supplemental Figure S4) in the K562 background and tested these cells’ resistance to Gal1-MMAF toxicity. As expected, sortilin KO cells are significantly less sensitive to Gal1-MMAF treatment than wild-type (WT) K562 cells. Furthermore, sortilin KO had no impact on the toxicity of MMAF alone, indicating that the effect was specific to Gal1 internalization (Figure 3a).

**Figure 3:**
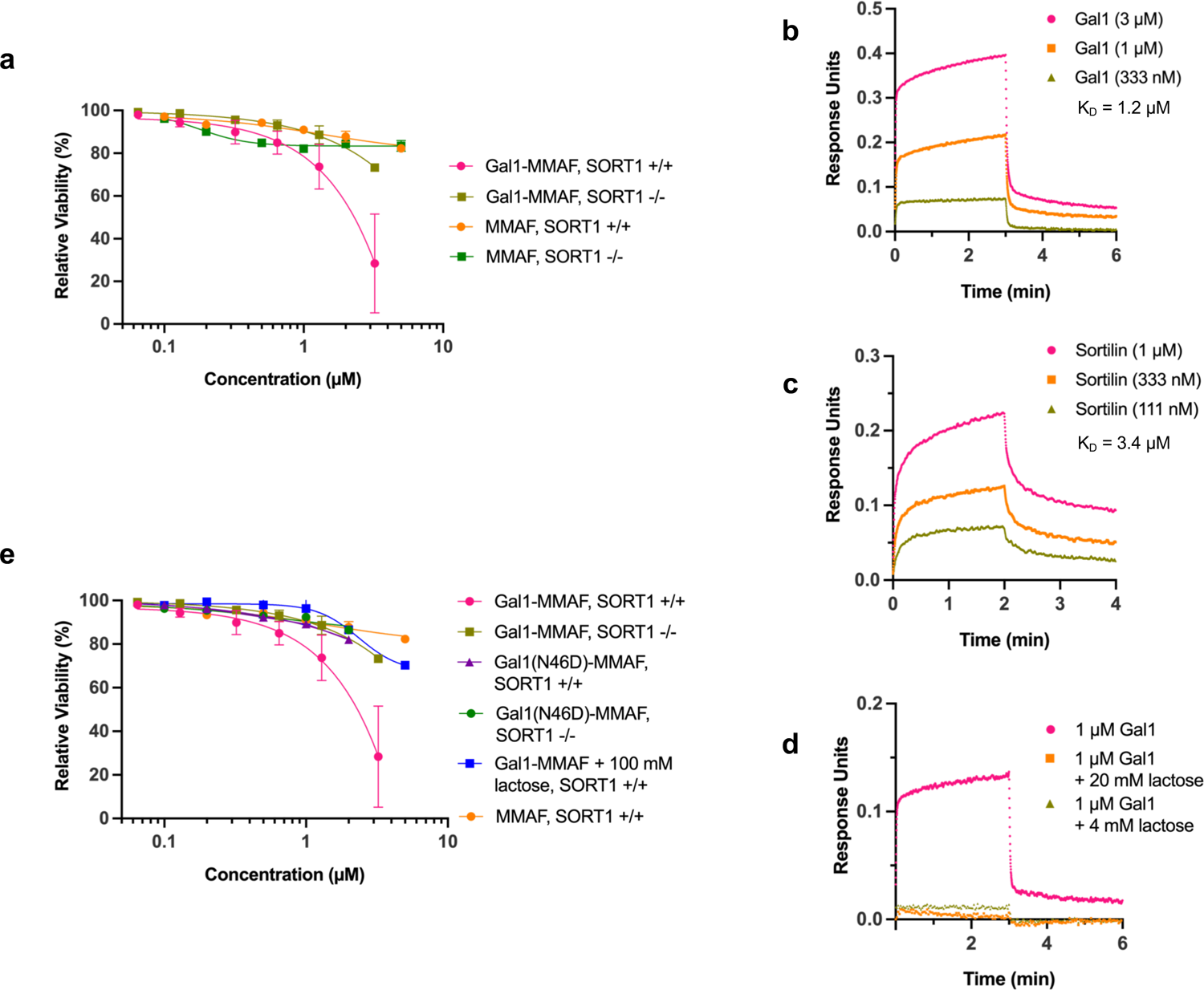
Sortilin Mediates Gal1-MMAF Toxicity via a Direct, Glycan-Dependent Interaction. **(A)** Flow cytometry ysis of WT and sortilin KO K562 cells treated with Gal1-MMAF or MMAF and stained with SYTOX to measure cell ility. **(B)** BLI analysis (Octet) of Gal1 binding to biotinylated sortilin immobilized on a streptavidin tip. erved K_D_ = ∼1.2 µM. **(C)** BLI analysis of sortilin binding to biotinylated Gal1 immobilized on a streptavidin BLI tip. erved K_D_ = ∼3.4 µM. **(D)** BLI analysis of Gal1 binding to biotinylated sortilin immobilized on a streptavidin BLI tip in presence or absence of lactose. **(E)** Flow cytometry analysis of WT and sortilin KO K562 cells treated with (N46D)-MMAF or Gal1-MMAF + 100 mM lactose and stained with SYTOX to measure cell viability.

We next investigated whether sortilin’s regulation of Gal1 internalization was mediated through a direct interaction. To test this, we analyzed the Gal1/sortilin interaction *in vitro* via bio-layer interferometry (BLI), a method for studying biomolecular interactions similar to surface plasmon resonance. We directly measured a specific interaction between sortilin and Gal1 with unit micromolar affinity, regardless of which protein was immobilized on the BLI tip (Figure 3b-c). The high on- and off-rates and micromolar binding affinity exhibited by the Gal1/sortilin interaction is typical of known protein-protein interactions at the cell surface^35^. Interestingly, this interaction appears to be significantly weaker than sortilin’s interaction with its canonical ligand, neurotensin, which is reported to bind sortilin with sub-nanomolar affinity^36^.

We also wished to test whether the interaction between Gal1 and sortilin was glycan-dependent. We investigated this by adding lactose, which is known to competitively inhibit Gal1’s glycan-binding activity^37,38^, to the BLI analyte solution. Addition of lactose completely abolished Gal1/sortilin binding (Figure 3d), indicating that glycans were involved in the Gal1/sortilin interaction. Based on these data, we created an MMAF conjugate using a Gal1 point mutant, Gal1(N46D), to test whether its toxicity was impacted. Gal1(N46D) has previously been shown to lack glycan-binding activity^39^, and we confirmed this by analyzing our construct via lactose affinity chromatography (Supplemental Figure S5). As expected, we observed that Gal1(N46D)-MMAF exhibited attenuated toxicity, similar to Gal1-MMAF in sortilin KO cells. Further, Gal1(N46D)-MMAF toxicity was not impacted by sortilin knockout (Figure 3e). Taken together, these data indicate that Gal1-MMAF toxicity is mediated by a glycan-dependent interaction with sortilin.

### Sortilin Mediates Lysosomal Trafficking of Gal1

We next wanted to measure the impact of sortilin on Gal1’s overall internalization, hypothesizing that sortilin KO would also result in reduced intracellular trafficking. To our surprise, however, when we treated sortilin KO cells with Gal1-AF647, we observed increased AF647 fluorescence in sortilin KO cells relative to WT (Figure 4a-c). To address this apparent discrepancy, these data led us to hypothesize that Gal1 may follow different intracellular trafficking pathways when internalized by sortilin KO vs WT K562 cells. Because not all of these pathways necessarily lead to Gal1-MMAF toxicity, which requires MMAF binding to tubulin in the cytosol^30^, this model could explain sortilin’s differential effects on toxicity and fluorescence.

**Figure 4:**
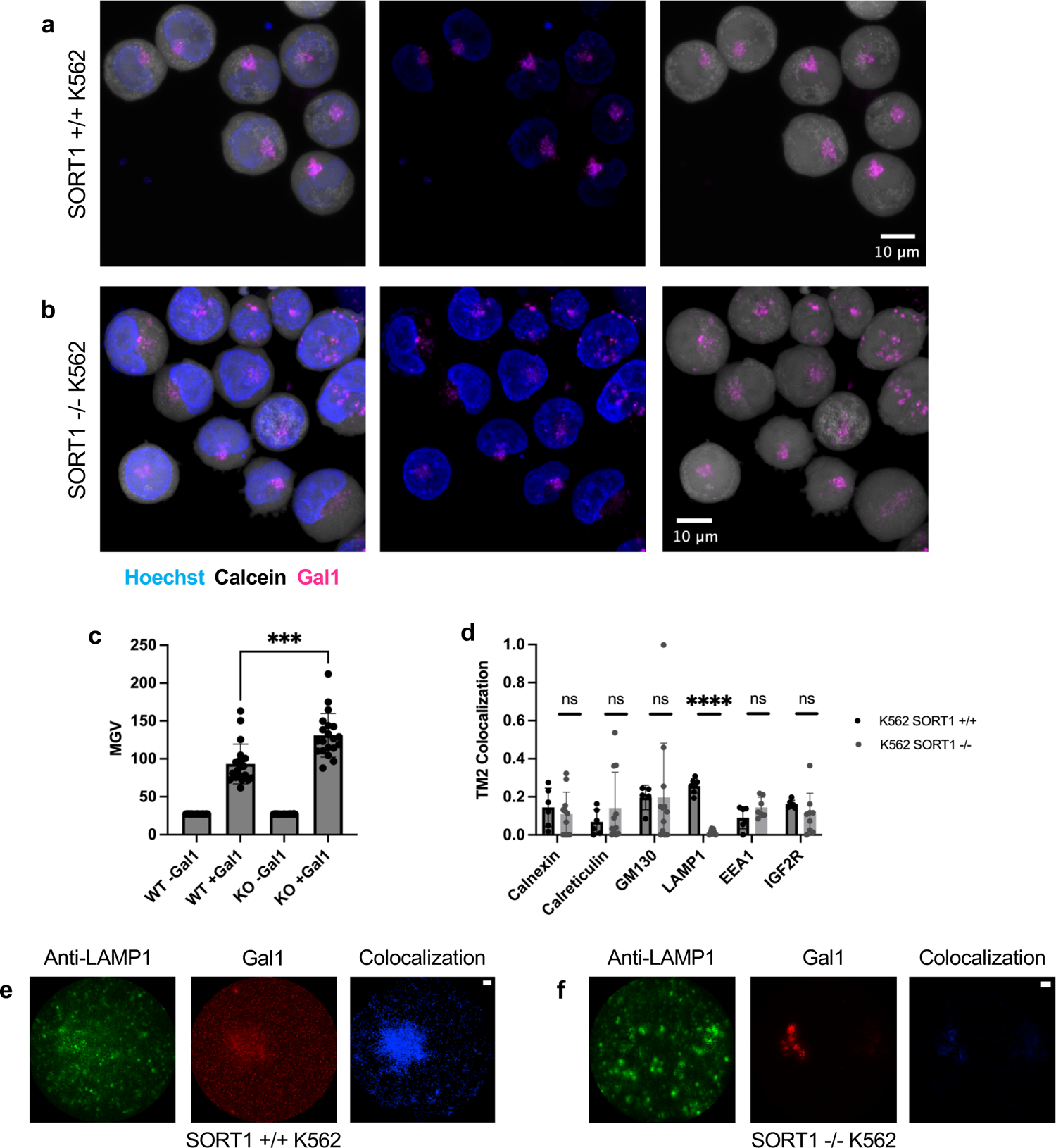
Sortilin is Required for Gal1 Trafficking to the Lysosome in K562 Cells. **(A, B)** Confocal images of live (A) and sortilin KO (B) K562 cells treated with Gal1-AF647 (2 µM, magenta) and stained with Hoechst (blue) and ein AM (white). **(C)** Fluorescence quantification of (A, B). *** = p <0.001. MGV = mean grey value. **(D)** WT and lin KO K562 cells were treated with Gal1-AF647 (5 µM) and stained with Hoechst and with one of six antibodies nst different organelle markers. Colocalization with each organelle marker was quantified using the thresholded Manders (TM2) algorithm. **** = p <1*10^-6^. ns = not significant (p > 0.05). **(E-F)** Representative images of WT (E) and lin KO (F) K562 cells stained with anti-LAMP1 (*green*), Gal1-AF647 (*red*). Colocalization mask (*blue*) also shown. e bar = 1 µm.

To explore this hypothesis, we used confocal immunofluorescence microscopy to investigate the subcellular localization of Gal1-AF647 in WT and sortilin KO K562 cells by analyzing its colocalization with a series of organelle markers for the ER, Golgi, and endolysosomal system. To quantify colocalization, we used the thresholded Manders coefficient (TM2), which quantified the fraction total Gal1-AF647 signal that colocalized with marker signal above an algorithmically-determined threshold^40^. Gal1-AF647 was observed to exhibit colocalization with all of these markers in WT K562 cells. Interestingly, sortilin KO resulted in a dramatic reduction in Gal1-AF647 colocalization with LAMP1, the marker for the lysosome, but did not significantly alter colocalization with any of the other markers (Figure 4d-f). These data provide a compelling explanation for why sortilin KO has opposing effects on Gal1-MMAF toxicity and Gal1-AF647 fluorescence. Due to the fact that Gal1 was conjugated to MMAF by a non-cleavable linker, digestion in the lysosome is likely necessary to enable MMAF’s toxicity by releasing it from Gal1. Further, these data indicate that sortilin is necessary for the trafficking of Gal1 to the lysosome.

### Gal1 Can Deliver Protein and RNA to the Cell Interior

The highly specific role of sortilin mediating trafficking of Gal1 to the lysosome presents an opportunity to use this system to direct other cargoes to this organelle, an application which is of significant interest to researchers and drug developers. As a proof-of-concept, we tested whether Gal1 can be used to deliver both model protein and model RNA cargoes.

We first fused Gal1 to GFP and tested its ability to be internalized by K562 and MRC5 cells. Flow cytometry and confocal imaging confirmed that Gal1-GFP was indeed internalized by both cell types (Figure 5a-c), indicating that Gal1 can be used to deliver proteins to the intracellular space. Additionally, while some methods for intracellular delivery of biomolecules such as cell-penetrating peptides can cause membrane permeabilization, which can result in toxicity and non-specific protein uptake^41^, we tested co-administration of Gal1 and GFP and observed no significant GFP uptake, indicating that the Gal1 internalization process does not affect the integrity of the plasma membrane (Figure 5d).

**Figure 5:**
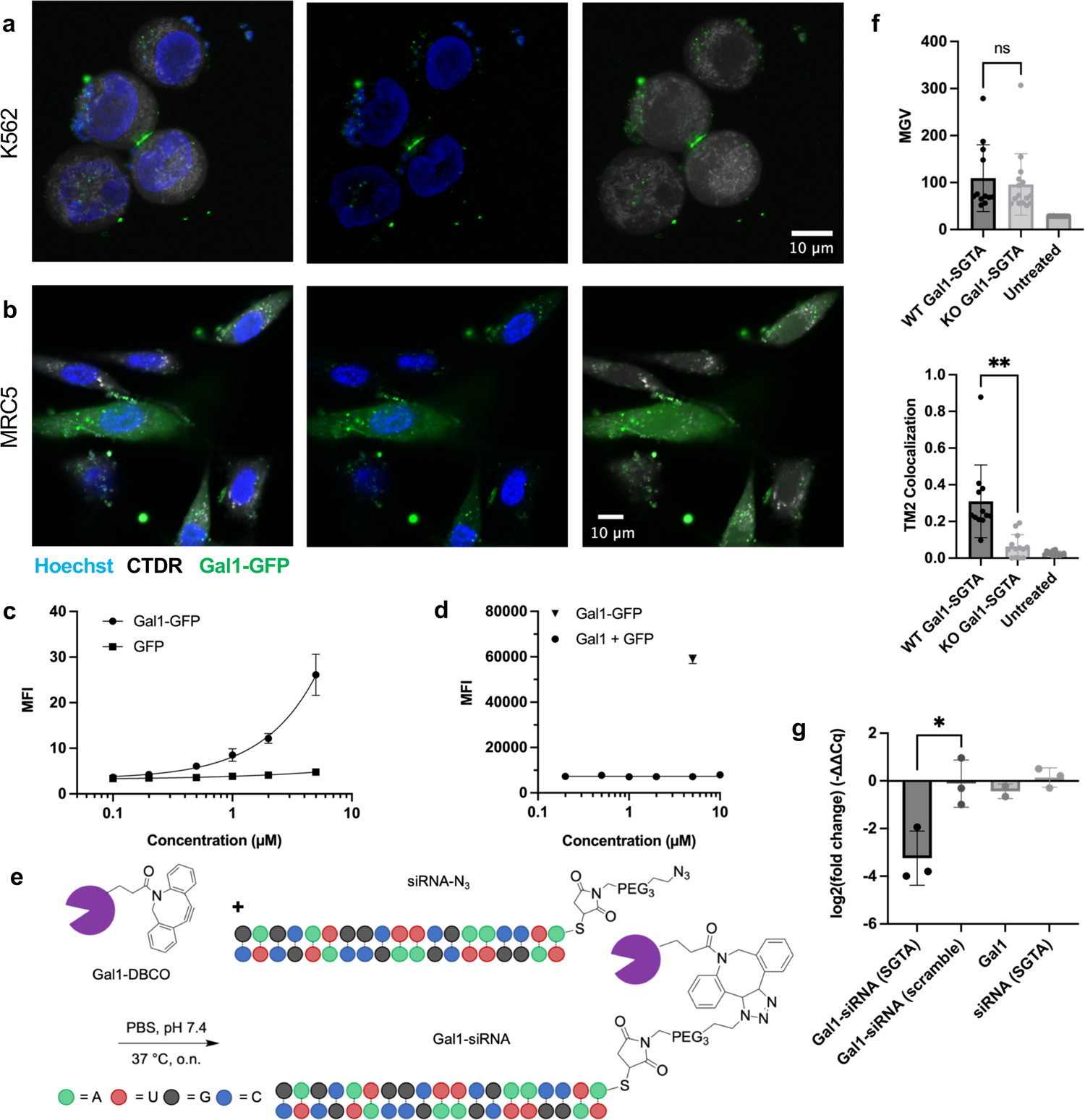
Gal1 Can Serve as a Delivery Vehicle for Protein or siRNA Cargo. **(A)** Confocal images of live K562 cells ed with Gal1-GFP (5 µM, green) and stained with Hoechst (blue) and CellTracker Deep Red (CTDR, white). **(B)** focal images of live MRC5 cells treated with Gal1-GFP (5 µM, green) and stained with Hoechst (blue) and CTDR te). **(C)** Flow cytometry analysis of live K562s treated with Gal1-GFP or GFP, then washed with 100 mM lactose re analysis. MFI = mean fluorescence intensity. **(D)** Flow cytometry analysis of live K562 cells treated with 5 µM and a concentration series of Gal1. Live K562 cells treated with 5 µM Gal1-GFP were used as a positive control. s were washed with 100 mM lactose before analysis. **(E)** Schematic of Gal1-siRNA preparation via click chemistry onjugation. DBCO-functionalized Gal1 (PEG_4_ linker not shown) and azide-functionalized siRNA were combined to Gal1-siRNA conjugates. **(F)** Fluorescence quantification (*top*) and LAMP1 colocalization analysis (*bottom*) of live and sortilin KO K562 cells treated with AF647-labeled Gal1-siRNA conjugates and then stained with Hoechst and LAMP1 before confocal imaging. MGV = mean grey value. ns = Not significant. ** = p <0.01. **(G)** qPCR analysis of 2 cells treated with a Gal1-siRNA conjugate targeting the housekeeping gene *SGTA*. * = p <0.05.

We then extended our investigation to siRNAs, which are of significant interest to drug developers due to their ability to knock down gene expression *in vivo*. We conjugated Gal1 to a model siRNA via click chemistry and characterized the resulting Gal1-siRNA conjugates (Figure 5e, Supplemental Figure S6). Gal1-siRNA conjugates functionalized with AF647 are internalized by K562 cells and trafficked to the lysosome in a sortilin-dependent manner, similarly to Gal1-AF647 (Figure 5f). Lastly, we also tested the ability of a prototypical Gal1-siRNA to knock-down a target gene using qPCR. Our results showed that Gal1-siRNA targeting the endogenous housekeeping gene *SGTA* effectively knocked down the target gene (Figure 5g), but Gal1 alone, naked *SGTA*-targeting siRNA, and Gal1 conjugated to a scrambled siRNA sequence did not exhibit this effect. These findings indicate that Gal1 can also be used as a delivery system for siRNAs similar to the ASGPR-targeting tri-GalNAc moiety currently used on clinically-approved siRNAs^42^. They also suggest that Gal1 has potential to serve as a platform for the development of new biologic therapies.

## Discussion

The work presented here indicates that Gal1 can follow multiple intracellular trafficking pathways once it is internalized into the endomembrane system. While studies of Gal1’s lectin activity typically explore the question of how its glycan-binding properties regulate the behavior of glycoprotein targets^12,14,43^, our research clearly demonstrates that glycoprotein binding partners also play an important role in regulating Gal1’s subcellular distribution. It is conceivable that the level of expression of sortilin, as well as the specific glycoforms it is displaying, combine to regulate the lysosomal trafficking and degradation of Gal1. Given the apparent importance of Gal1 expression levels to key disease processes, including cancer progression and immune activation^10,14,19^, this relationship deserves further study.

Our discovery that sortilin is involved in the lysosomal trafficking of Gal1 fits into existing knowledge about sortilin’s functions. For example, sortilin is known to mediate lysosomal trafficking of sphingolipid activator proteins from the Golgi^32^ and transport of β-site β-amyloid precursor protein-cleaving enzyme 1 (BACE1) from the endolysosomal system to the trans-Golgi network (TGN)^33^. Interestingly, BACE1 trafficked to the TGN can be further retrograde trafficked to the ER^33^, a known mechanism of cytosolic trafficking of extracellular proteins^28,44^.

This work also further supports the notion that lectins have specificity not only to certain glycans, but also to specific glycoproteins or glycolipids to which those glycans are conjugated. For example, researchers had long believed that the lectin Siglec-7 bound to a broad variety of ligands bearing sialoside glycans, as suggested by glycan array data^45^. However, Wisnovsky and colleagues used a CRISPR screen to identify Siglec-7 ligands. In the context of an entire cell, Siglec-7 was revealed to have a professional ligand: the sialomucin CD43^46^. Another example, P-selectin, was originally thought to bind sialyl-Lewis X structures, but its binding is almost entirely determined by P-selectin glycoprotein ligand-1 (PSGL-1)^47^. Similarly, based on glycan array experiments, Gal1 has been shown to prefer the unsialylated LacNAc motifs commonly found on N-glycans^45^. These motifs are ubiquitous on glycoproteins, yet knockout of sortilin completely abolishes Gal1 lysosomal trafficking, which requires a specific interaction with the sortilin glycoprotein.

An intriguing disease-relevant question we have not yet answered involves how extracellular Gal1 reaches the nucleus, which has been demonstrated in previous work^17,22^. Interestingly, some proteins such as the transcription factor ATF6 and bacterial toxins like Ricin and Shiga toxin have been reported to undergo retrograde trafficking through the Golgi and ER to the nucleus or cytosol^28,44^. Another study highlights the existence of “nuclear-associated endosomes” (NAEs) and finds that pseudomonas exotoxin A can traffic to NAEs in a Golgi-independent fashion. These NAEs can fuse with the nuclear envelope, resulting in transport of their contents to the nucleoplasm^48^. Further investigation will be required shed light whether Gal1 reaches the nucleus through one of these pathways or via a novel and entirely uncharacterized one. The insight that sortilin is involved in the transport of Gal1 to the lysosome raises the question of whether the sortilin-Gal1 axis can regulate the surface lifetime and intracellular trafficking of cell-surface receptors. It is well-established that changes in the glycosylation of cell surface receptors such as Her2 and CD22/Siglec-2 can affect their rate of internalization and intracellular trafficking, but the mechanism of this regulation is unclear^29^. It is conceivable that Gal1 could act as an adaptor protein on the sortilin receptor, enabling it to regulate trafficking of diverse glycoprotein targets.

While our prototypical LDC demonstrated the utility of this class of molecules as research tools with applications to functional genomics, we can also envision its potential future applicability as therapeutics that selectively target cells based on the glycan-dependent interactions of lectins. Gal1 ligands are upregulated in cancers including glioblastoma, T-cell lymphomas, and pancreatic adenocarcinoma^49,50^, making our Gal1-MMAF prototype an excellent candidate for a potential therapeutic LDC. Notably, several bispecific molecules that form ternary interactions with a lysosomal trafficking receptor and a cell-surface protein of interest can also function as LYTACs, effecting degradation of the target protein^51^. The Gal1/sortilin interaction could also conceivably play such a lysosomal targeting role.

Lastly, lysosomal delivery of biomolecule cargoes is of significant interest to drug developers. Many FDA-approved siRNA therapeutics rely on the asialoglycoprotein receptor (ASGPR) to deliver siRNAs to the lysosomes of hepatocytes^52^. To date, only liver targets have been successfully drugged with this modality^42^. The robust trafficking of Gal1 to the lysosome in sortilin-expressing cells raises the possibility of delivering therapeutic siRNAs to targets outside the liver, particularly to the brain, where sortilin is highly expressed^53^.

## Supporting Information

Additional experimental details, materials, methods, spectra, and characterization data can be found in the Supporting Information.

## Acknowledgements

J.D. would like to thank Dr. Joseph A. Buonomo, Dr. Ryan A. Flynn, Dr. Rishikesh U. Kulkarni, Dr. Marie A. Hollenhorst, Dr. Lindsay E. Guzmán, and Dr. Kevin J. Bruemmer for useful discussions. J.D. would also like to thank Dr. Shaogeng “Steven” Tang for technical assistance with BLI experiments and analysis and Sophia Shi for assistance with colocalization data analysis. Some figures were created using BioRender.

## Experimental Methods

### Protein & RNA Conjugation and Click Chemistry

NHS ester-based conjugations to protein were carried out in PBS + 150 mM NaCl + 5 mM TCEP at pH 8.3. Buffer for functionalization of Gal1 was supplemented with 10 mM lactose. Reactions were carried out overnight at r.t. or 37 °C with 10 molar equivalent NHS ester reagent (with the exception of NHS-based biotinylation, which was carried out with 3 molar eq. at 37 °C for 1 hr). Maleimide-based functionalization of thiol-modified siRNAs (custom-ordered from Millipore-Sigma) was accomplished in PBS + 5 mM TCEP at pH 7.3, reacted overnight at 37 °C with 10 molar equivalent maleimide reagent. Copper-free click reactions between DBCO- and azide-functionalized molecules were performed in PBS + 5 mM TCEP at pH 7-7.5 at r.t. or 37 °C overnight with a mole ratio between 1-3. Buffer exchange and removal of excess conjugation reagent was performed using Zeba columns (Fisher Scientific, Cat. #89890, 89891, 89882, 89893) according to the manufacturer protocol.

### Top-Down Protein Mass Spectrometry

After protein expression and purification, the protein samples were further buffer exchanged using a methanol/chloroform/water precipitation and resolubilization method^54^. Here, 300 µL of 10 mM TCEP prepared in cold LC/MS-grade water (4 °C) was added to 100 µL of protein solution. Then, 400 µL of cold methanol (−20 °C) was added to the protein solution and vortexed for 30 sec followed by 100 µL of cold chloroform (−20 °C) and an additional 30 sec of vortexing. The sample was centrifuged for 10 min at 18,000 xG at 4 °C after which a biphasic mixture was created with a protein pellet present at the interface. The top layer of the solution was discarded without disturbing the protein pellet. Then, 400 µL of cold methanol (−20 °C) was added to the sample and gently vortexed. The sample was centrifuged for 10 min at 18,000 xG at 4 °C after which the supernatant was discarded. The cold methanol and centrifugation washing step was repeated two additional times. Protein pellets were resolubilized with 4 µL of 80% formic acid (−20 °C) and diluted to 1% formic acid with 80:20 water:acetonitrile. Top-down LC-MS was carried out using an Agilent 1260 Infinity II high performance liquid chromatography (HPLC) system coupled to an Agilent 6230 ToF LC/MS (Agilent Technologies). Samples were injected onto an Agilent PLRP-S column (2.1 × 50 mm, 5 µm particle size, 1,000 Å pore size) using a gradient of 10% to 90% mobile phase B (0–5 min, 10% B; 5–15 min, 20%–65% B; 15–18 min, 65%–90% B, 18–22 min, 90% B; 22–25 min, 10% B; mobile phase A set to 0.2% formic acid in water; mobile phase B set to 0.2% formic acid in acetonitrile). Flow rate was set to 200 µL/min with a column temperature of 60 °C. Mass spectra were taken at a scan rate of 1 Hz over a 200–3200 m/z scan range with the ToF set to extended dynamic range. The mass spectrometer was operated using a dual Agilent jet stream (AJS) high-sensitivity ion source with the following instrument parameters: gas temperature (275 °C), drying gas (12 L/min), nebulizer (40 psi), sheath gas temperature (400 °C), sheath gas flow (12 L/min), VCap (3000 V), nozzle voltage (2000 V), fragmentor (250 V), skimmer (65 V), and Oct 1 RF Vpp (750 V). Mass spectra were output from the MassHunter (Agilent Technologies) software and analyzed using MASH Native^55^ and UniDec^56^.

### Mammalian Tissue Culture & Generation of Knockout Cells

All cell lines were maintained in DMEM (Thermo Scientific, Cat. #11965118) + 10% fetal bovine serum (FBS, various suppliers) + 1X PenStrep (Fisher Scientific, Cat. #sv30010) or 1X Gibco antibiotic-antimycotic (Fisher Scientific, Cat. #15-240-096) at 37 °C and 5% CO_2_ in a humidified incubator. Cell quantity and viability were determined via staining by Trypan Blue (VWR, Cat. #VWRVK940). Sortilin KO cells were generated using a Lonza 4D Nucleofector X nucleofector and a Synthego v2 gene KO kit with *SORT1* sgRNA (custom-ordered) according to the manufacturer protocol.

### Gal1 Internalization Assay, Confocal Fluorescence Microscopy

Cells were plated in borosilicate 1.5 chambered coverglass plates (Thomas Scientific, Cat. #9384V15) and treated with Gal1-AF647 or Gal1-GFP overnight. For K562 cells, plates were pretreated with 10 µg/mL human fibronectin (ThermoFisher Scientific, Cat. #33-016-015) at 37 °C for ≥1 hr. For live cell imaging, cells were washed twice with ice-cold PBS + 100 mM lactose before staining with 20 µM Hoechst 33342 (ThermoFisher Scientific, Cat. #62249) and 1 µg/mL Calcein-AM (for Gal1-AF647, BioLegend, Cat. #425201) or 1 µM CellTracker Deep Red (for Gal1-GFP, Thermo Scientific, Cat. #C34565) for 15 min at 4 °C. Dye solution was removed and replaced with PBS for imaging. For immunofluorescence, cells were fixed in 4% paraformaldehyde (Alfa Aesar, Cat. #J61899-AP) for 15 min at r.t. and permeabilized with 0.1% Triton X-100 for 10 min at r.t. After blocking with 4% goat serum + 1% BSA + 0.05% Tween-20, cells were stained with primary antibody (1:1,000 in PBS + 1% BSA + 0.05% Tween-20) for 1 hr at r.t. followed by secondary antibody (1:1,000 in PBS + 1% BSA + 0.05% Tween-20) with 20 µM Hoechst for 1 hr at r.t. Cells were imaged using a Nikon Eclipse Ti confocal microscope with a 60X oil-immersion objective. 10-slice Z-stacks of individual cells were analyzed in ImageJ for fluorescence intensity (mean grey value, MGV) and colocalization (thresholded Manders coefficient, tM) where appropriate^40^.

### Gal1 Internalization and Toxicity Assays, Flow Cytometry

Cells were plated in 96-well plates and treated with purified Gal1 constructs overnight, or 48 hr for Gal1-MMAF conjugates. Cells were then washed twice (with centrifugation at 500-600 xG) with ice cold PBS + 100 mM lactose (for experiments measuring Gal1 fluorescence) or ice-cold PBS. Cells were then stained with SYTOX Red (for Gal1-GFP, ThermoFisher Scientific, Cat. #S34859) or SYTOX Green (all other experiments, ThermoFisher Scientific, Cat. #S7020) at 4 °C according to the manufacturer protocol. In SYTOX assays, positive control samples for cell death were prepared by heating cells to 95 °C for 10 min while experimental cell samples were maintained on ice. Cells were analyzed using a Miltenyi MACSQuant Analyzer 10 flow cytometer.

### CRISPR Screen

K562 cells transduced with a previously-described 200,000 sgRNA genome-wide CRISPRi library^57^ were treated with 1 µM Gal1-MMAF for 48 hr. Cells were then washed 2x with PBS (with centrifugation at 350 xG) to remove excess toxin and dead cells. Cells were counted to determine % selection and coverage. Selection treatment was repeated two additional times. Cells were maintained at ≥500x guide coverage throughout the experiment. After treatment, genomic DNA (gDNA) was prepared in a plasmid- and PCR product-free work area using a QIAamp DNA Blood Maxi Kit (Qiagen, Cat. #51192) according to manufacturer instructions and previously published protocols^27^. sgRNA-encoding genomic loci were amplified from up to 400 µg gDNA via PCR using a Herculase II Fusion Polymerase PCR kit (Agilent, Cat. #600677). Pooled reaction products from each treatment group were uniquely barcoded in a subsequent PCR reaction followed by electrophoresis on 2% TBE-agarose gel at 120 V. Amplicons were purified via a Zymoclean Gel DNA Recovery kit (Zymo Research, Cat. #D4007) according to the manufacturer protocol before Qubit quantification and Illumina NextSeq sequencing at 250x coverage as previously described^29^. Sequencing data were analyzed using the casTLE algorithm^27^.

### Binding Assays

Binding assays were carried out using an Octet BLI system as previously described^58^. An Octet streptavidin (SA) biosensor tip (Sartorius, Cat. #18-5019) was used to immobilize biotinylated Gal1 or sortilin before measurement of analyte association.

### Quantitative PCR

Cells were treated with Gal1-siRNA conjugates for 48 hr. RNA was extracted and purified via a Trizol (ThermoFisher Scientific, Cat. #15596026) extraction. Cell pellets were resuspended in 100 µL PBS before addition of 1 mL of Trizol with thorough mixing. After 5 min incubation at r.t. in Trizol, 200 µL chloroform was added and incubated for an additional 2 min. Samples were then centrifuged at 12,000 xG for 2 min to partition solvents. The aqueous (top) layer was recovered and 3x recovered volume of neat ethanol was added. After thorough mixing, the entire volume of each sample was loaded onto a Zymogen Zymo-Spin IIN column (Zymo Research, Cat. #C1019) and spun at ≥15,000 xG for 1 min, repeating and discarding flow-through until the entire sample had been loaded. Columns were then washed twice with 300 µL 80% ethanol, followed by an additional dry spin to fully clear the membrane. Samples were eluted with 30 µL nuclease-free water at 55-65 °C, incubating 5 min before the final spin to recover purified RNA. Purified RNA was quantified via absorbance at 260 nm and concentrations were normalized across samples via addition of nuclease-free water. Quantitative PCR (qPCR) reactions were prepared using an Invitrogen Superscript III Platinum SYBR Green One-Step qRT-PCR kit (Fisher Scientific, Cat. #11736051) according to the manufacturer protocol and analyzed on a BioRad CFX96 real-time qPCR system.

## References

1. Modenutti, C. P., Capurro, J. I. B., Di Lella, S., & Martí, M. A. (2019). The Structural Biology of Galectin-Ligand Recognition: Current Advances in Modeling Tools, Protein Engineering, and Inhibitor Design. Frontiers in Chemistry, 7(December). 10.3389/fchem.2019.00823

2. Leppänen, A., Stowell, S., Blixt, O., & Cummings, R. D. (2005). Dimeric galectin-1 binds with high affinity to α2,3-sialylated and non-sialylated terminal N-acetyllactosamine units on surface-bound extended glycans. Journal of Biological Chemistry, 280(7), 5549–5562. 10.1074/jbc.M412019200

3. Camby, I., Le Mercier, M., Lefranc, F., & Kiss, R. (2006). REVIEW Galectin-1: a small protein with major functions. Glycobiology, 16(11), 137–157. 10.1093/glycob/cwl025

4. Astorgues-Xerri, L., Riveiro, M. E., Tijeras-Raballand, A., Serova, M., Neuzillet, C., Albert, S., Raymond, E., & Faivre, S. (2014). Unraveling galectin-1 as a novel therapeutic target for cancer. In Cancer Treatment Reviews (Vol. 40, Issue 2, pp. 307– 319). 10.1016/j.ctrv.2013.07.007

5. Cousin, J. M., & Cloninger, M. J. (2016). The role of galectin-1 in cancer progression, and synthetic multivalent systems for the study of Galectin-1. International Journal of Molecular Sciences, 17(9). 10.3390/ijms17091566

6. Hsu, Y. L., Wu, C. Y., Hung, J. Y., Lin, Y. S., Huang, M. S., & Kuo, P. L. (2013). Galectin-1 promotes lung cancer tumor metastasis by potentiating integrin α6β4 and Notch1/Jagged2 signaling pathway. Carcinogenesis, 34(6), 1370–1381. 10.1093/carcin/bgt040

7. Pace, K. E., Lee, C., Stewart, P. L., & Baum, L. G. (1999). Restricted Receptor Segregation into Membrane Microdomains Occurs on Human T Cells During Apoptosis Induced by Galectin-1. The Journal of Immunology, 163(7), 3801–3811. 10.4049/jimmunol.163.7.3801

8. Symons, A., Cooper, D. N., & Barclay, A. N. (2000). Characterization of the interaction between galectin-1 and lymphocyte glycoproteins CD45 and Thy-1. Glycobiology, 10(6), 559–563. 10.1093/glycob/10.6.559

9. Miller, M. C., Nesmelova, I. V, Platt, D., Klyosov, A., & Mayo, K. H. (2009). The carbohydrate-binding domain on galectin-1 is more extensive for a complex glycan than for simple saccharides: Implications for galectin-glycan interactions at the cell surface. Biochemical Journal, 421(2), 211–221. 10.1042/BJ20090265

10. Hernandez, J. D., Nguyen, J. T., He, J., Wang, W., Ardman, B., Green, J. M., Fukuda, M., & Baum, L. G. (2006). Galectin-1 Binds Different CD43 Glycoforms to Cluster CD43 and Regulate T Cell Death. The Journal of Immunology, 177(8), 5328–5336. 10.4049/jimmunol.177.8.5328

11. Belardi, B., Odonoghue, G. P., Smith, A. W., Groves, J. T., & Bertozzi, C. R. (2012). Investigating cell surface galectin-mediated cross-linking on glycoengineered cells. Journal of the American Chemical Society, 134(23), 9549–9552. 10.1021/ja301694s

12. Jeschke, U., Karsten, U., Wiest, I., Schulze, S., Kuhn, C., Friese, K., & Walzel, H. (2006). Binding of galectin-1 (gal-1) to the Thomsen-Friedenreich (TF) antigen on trophoblast cells and inhibition of proliferation of trophoblast tumor cells in vitro by gal-1 or an anti-TF antibody. Histochemistry and Cell Biology, 126(4), 437–444. 10.1007/s00418-006-0178-1

13. Rabinovich, G. A., Ariel, A., Hershkoviz, R., Hirabayashi, J., Kasai, K. I., & Lider, O. (1999). Specific inhibition of T-cell adhesion to extracellular matrix and proinflammatory cytokine secretion by human recombinant galectin-1. Immunology, 97(1), 100–106. 10.1046/j.1365-2567.1999.00746.x

14. Fulcher, J. A., Hashimi, S. T., Levroney, E. L., Pang, M., Gurney, K. B., Baum, L. G., & Lee, B. (2006). Have Enhanced Migration through Extracellular Matrix 1. Gene, 19.

15. Blaser, C., Kaufmann, M., Müller, C., Zimmermann, C., Wells, V., Mallucci, L., & Pircher, H. (1998). β-galactoside-binding protein secreted by activated T cells inhibits antigen-induced proliferation of T cells. European Journal of Immunology, 28(8), 2311– 2319. 10.1002/(SICI)1521-4141(199808)28:08<2311::AID-IMMU2311>3.0.CO;2-G

16. Rubinstein, N., Alvarez, M., Zwirner, N. W., Toscano, M. A., Ilarregui, J. M., Bravo, A., Mordoh, J., Fainboim, L., Podhajcer, O. L., & Rabinovich, G. A. (2004). Targeted inhibition of galectin-1 gene expression in tumor cells results in heightened T cell-mediated rejection: A potential mechanism of tumor-immune privilege. Cancer Cell, 5(3), 241–251. 10.1016/S1535-6108(04)00024-8

17. Bhat, R., Belardi, B., Mori, H., Kuo, P., Tam, A., Hines, W. C., Le, Q. T., Bertozzi, C. R., & Bissell, M. J. (2016). Nuclear repartitioning of galectin-1 by an extracellular glycan switch regulates mammary morphogenesis. Proceedings of the National Academy of Sciences of the United States of America, 113(33), E4820–E4827. 10.1073/pnas.1609135113

18. Sharanek, A., Burban, A., Hernandez-Corchado, A., Madrigal, A., Fatakdawala, I., Najafabadi, H. S., Soleimani, V. D., & Jahani-Asl, A. (2021). Transcriptional control of brain tumor stem cells by a carbohydrate binding protein. Cell Reports, 36(9). 10.1016/j.celrep.2021.109647

19. Gao, Y., Li, X., Shu, Z., Zhang, K., Xue, X., Li, W., Hao, Q., Wang, Z., Zhang, W., Wang, S., Zeng, C., Fan, D., Zhang, W., Zhang, Y., Zhao, H., Li, M., & Zhang, C. (2018). Nuclear galectin-1-FOXP3 interaction dampens the tumor-suppressive properties of FOXP3 in breast cancer. Cell Death and Disease, 9(4). 10.1038/s41419-018-0448-6

20. Voss, P. G., Gray, R. M., Dickey, S. W., Wang, W., Park, J. W., Kasai, K. ichi, Hirabayashi, J., Patterson, R. J., & Wang, J. L. (2008). Dissociation of the carbohydrate-binding and splicing activities of galectin-1. Archives of Biochemistry and Biophysics, 478(1), 18–25. 10.1016/j.abb.2008.07.003

21. Vyakarnam, A., Dagher, S. F., Wang, J. L., & Patterson, R. J. (1997). Evidence for a Role for Galectin-1 in Pre-mRNA Splicing. Molecular and Cellular Biology, 17(8), 4730–4737. 10.1128/mcb.17.8.4730

22. Park, J. W., Voss, P. G., Grabski, S., Wang, J. L., & Patterson, R. J. (2001). Association of galectin-1 and galectin-3 with Gemin4 in complexes containing the SMN protein. Nucleic Acids Research, 29(17), 3595–3602. 10.1093/nar/29.17.3595

23. Wei, J., Li, D. K., Hu, X., Cheng, C., & Zhang, Y. (2021). Galectin-1–RNA interaction map reveals potential regulatory roles in angiogenesis. FEBS Letters, 595(5), 623–636. 10.1002/1873-3468.14047

24. Thijssen, V. L., Barkan, B., Shoji, H., Aries, I. M., Mathieu, V., Deltour, L., Hackeng, T. M., Kiss, R., Kloog, Y., Poirier, F., & Griffioen, A. W. (2010). Tumor cells secrete galectin-1 to enhance endothelial cell activity. Cancer Research, 70(15), 6216–6224. 10.1158/0008-5472.CAN-09-4150

25. Cho, M., & Cummings, R. D. (1995). Galectin-1, a β-galactoside-binding lectin in Chinese hamster ovary cells. I. Physical and chemical characterization. Journal of Biological Chemistry, 270(10), 5198–5206. 10.1074/jbc.270.10.5198

26. Davuluri, G. V. N., Chen, C. C., Chiu, Y. C., Tsai, H. W., Chiu, H. C., Chen, Y. L., Tsai, P. J., Kuo, W. T., Tsao, N., Lin, Y. S., & Chang, C. P. (2021). Autophagy Drives Galectin-1 Secretion From Tumor-Associated Macrophages Facilitating Hepatocellular Carcinoma Progression. Frontiers in Cell and Developmental Biology, 9(September), 1–15. 10.3389/fcell.2021.741820

27. Morgens, D. W., Deans, R. M., Li, A., & Bassik, M. C. (2016). Systematic comparison of CRISPR/Cas9 and RNAi screens for essential genes. Nature Biotechnology, 34(6), 634– 636. 10.1038/nbt.3567

28. Bassik, M. C., Kampmann, M., Lebbink, R. J., Wang, S., Hein, M. Y., Poser, I., Weibezahn, J., Horlbeck, M. A., Chen, S., Mann, M., Hyman, A. A., Leproust, E. M., McManus, M. T., & Weissman, J. S. (2013). A systematic mammalian genetic interaction map reveals pathways underlying ricin susceptibility. Cell, 152(4), 909–922. 10.1016/j.cell.2013.01.030

29. Tsui, C. K., Barfield, R. M., Fischer, C. R., Morgens, D. W., Li, A., Smith, B. A. H., Gray, M. A., Bertozzi, C. R., Rabuka, D., & Bassik, M. C. (2019). CRISPR-Cas9 screens identify regulators of antibody–drug conjugate toxicity. Nature Chemical Biology, 15(10), 949–958. 10.1038/s41589-019-0342-2

30. Hingorani, D. V, Doan, M. K., Camargo, M. F., Aguilera, J., Song, S. M., Pizzo, D., Scanderbeg, D. J., Cohen, E. E. W., Lowy, A. M., Adams, S. R., & Advani, S. J. (2020). Precision Chemoradiotherapy for HER2 Tumors Using Antibody Conjugates of an Auristatin Derivative with Reduced Cell Permeability. Molecular Cancer Therapeutics, 19(1), 157–167. 10.1158/1535-7163.MCT-18-1302

31. Jewett, J. C., Sletten, E. M., & Bertozzi, C. R. (2010). Rapid Cu-free click chemistry with readily synthesized biarylazacyclooctynones. Journal of the American Chemical Society, 132(11), 3688–3690. 10.1021/ja100014q

32. Lefrancois, S., Zeng, J., Hassan, A. J., Canuel, M., & Morales, C. R. (2003). The lysosomal trafficking of sphingolipid activator proteins (SAPs) is mediated by sortilin. EMBO Journal, 22(24), 6430–6437. 10.1093/emboj/cdg629

33. Finan, G. M., Okada, H., & Kim, T. W. (2011). BACE1 retrograde trafficking is uniquely regulated by the cytoplasmic domain of sortilin. Journal of Biological Chemistry, 286(14), 12602–12616. 10.1074/jbc.M110.170217

34. Richner, M., Pallesen, L. T., Ulrichsen, M., Poulsen, E. T., Holm, T. H., Login, H., Castonguay, A., Lorenzo, L. E., Gonçalves, N. P., Andersen, O. M., Lykke-Hartmann, K., Enghild, J. J., Rønn, L. C. B., Malik, I. J., De Koninck, Y., Bjerrum, O. J., Vægter, C. B., & Nykjær, A. (2019). Sortilin gates neurotensin and BDNF signaling to control peripheral neuropathic pain. Science Advances, 5(6), 9946–9965. 10.1126/sciadv.aav9946

35. Chen, J., Sawyer, N., & Regan, L. (2013). Protein-protein interactions: General trends in the relationship between binding affinity and interfacial buried surface area. Protein Science, 22(4), 510–515. 10.1002/pro.2230

36. Mazella, J., Zsürger, N., Navarro, V., Chabry, J., Kaghad, M., Caput, D., Ferrara, P., Vita, N., Gully, D., Maffrand, J. P., & Vincent, J. P. (1998). The 100-kDa neurotensin receptor is gp95/sortilin, a non-G-protein-coupled receptor. Journal of Biological Chemistry, 273(41), 26273–26276. 10.1074/jbc.273.41.26273

37. Rabinovich, G. A., Alonso, C. R., Sotomayor, C. E., Durand, S., Bocco, J. L., & Riera, C. M. (2000). Molecular mechanisms implicated in galectin-1-induced apoptosis: Activation of the AP-1 transcription factor and downregulation of Bcl-2. Cell Death and Differentiation, 7(8), 747–753. 10.1038/sj.cdd.4400708

38. Amano, M., Galvan, M., He, J., & Baum, L. G. (2003). The ST6Gal I sialyltransferase selectively modifies N-glycans on CD45 to negatively regulate galectin-1-induced CD45 clustering, phosphatase modulation, and T cell death. Journal of Biological Chemistry, 278(9), 7469–7475. 10.1074/jbc.M209595200

39. Hirabayashi, J., & Kasai, K. I. (1991). Effect of amino acid substitution by site-directed mutagenesis on the carbohydrate recognition and stability of human 14-kDa β-galactoside-binding lectin. Journal of Biological Chemistry, 266(35), 23648–23653.

40. Dunn, K. W., Kamocka, M. M., & McDonald, J. H. (2011). A practical guide to evaluating colocalization in biological microscopy. American Journal of Physiology. Cell Physiology, 300(4), C723–42. 10.1152/ajpcell.00462.2010

41. Yu, S., Yang, H., Li, T., Pan, H., Ren, S., Luo, G., Jiang, J., Yu, L., Chen, B., Zhang, Y., Wang, S., Tian, R., Zhang, T., Zhang, S., Chen, Y., Yuan, Q., Ge, S., Zhang, J., & Xia, N. (2021). Efficient intracellular delivery of proteins by a multifunctional chimaeric peptide in vitro and in vivo. Nature Communications, 12(1). 10.1038/s41467-021-25448-z

42. Hu, B., Zhong, L., Weng, Y., Peng, L., Huang, Y., Zhao, Y., & Liang, X. J. (2020). Therapeutic siRNA: state of the art. In Signal Transduction and Targeted Therapy (Vol. 5, Issue 1). 10.1038/s41392-020-0207-x

43. Starossom, S. C., Mascanfroni, I. D., Imitola, J., Cao, L., Raddassi, K., Hernandez, S. F., Bassil, R., Croci, D. O., Cerliani, J. P., Delacour, D., Wang, Y., Elyaman, W., Khoury, S. J., & Rabinovich, G. A. (2012). Galectin-1 Deactivates Classically Activated Microglia and Protects from Inflammation-Induced Neurodegeneration. Immunity, 37(2), 249–263. 10.1016/j.immuni.2012.05.023

44. Sandvig, K., Grimmer, S., Lauvrak, S., Torgersen, M., Skretting, G., Van Deurs, B., & Iversen, T. (2002). Pathways followed by ricin and Shiga toxin into cells. Histochemistry and Cell Biology, 117(2), 131–141. 10.1007/s00418-001-0346-2

45. Arthur, C. M., Rodrigues, L. C., Baruffi, M. D., Sullivan, H. C., Heimburg-Molinaro, J., Smith, D. F., Cummings, R. D., & Stowell, S. R. (2015). Examining Galectin Binding Specificity Using Glycan Microarrays. In S. R. Stowell & R. D. Cummings (Eds.), Methods in Molecular Biology (Vol. 1207, Issue 4, pp. 115–131). Springer New York. 10.1007/978-1-4939-1396-1_8

46. Wisnovsky, S., Möckl, L., Malaker, S. A., Pedram, K., Hess, G. T., Riley, N. M., Gray, M. A., Smith, B. A. H., Bassik, M. C., Moerner, W. E., & Bertozzi, C. R. (2021). Genome-wide CRISPR screens reveal a specific ligand for the glycan-binding immune checkpoint receptor Siglec-7. Proceedings of the National Academy of Sciences of the United States of America, 118(5). 10.1073/pnas.2015024118

47. Norgard, K. E., Moore, K. L., Diaz, S., Stults, N. L., Ushiyama, S., McEver, R. P., Cummings, R. D., & Varki, A. (1993). Characterization of a specific ligand for P-selectin on myeloid cells. A minor glycoprotein with sialylated O-linked oligosaccharides. Journal of Biological Chemistry, 268(17), 12764–12774. 10.1016/s0021-9258(18)31454-6

48. Chaumet, A., Wright, G. D., Seet, S. H., Tham, K. M., Gounko, N. V, & Bard, F. (2015). Nuclear envelope-associated endosomes deliver surface proteins to the nucleus. Nature Communications, 6. 10.1038/ncomms9218

49. Yang, W., Wu, P. fei, Ma, J. xing, Liao, M. jun, Wang, X. hui, Xu, L. shan, Xu, M. hui, & Yi, L. (2019). Sortilin promotes glioblastoma invasion and mesenchymal transition through GSK-3β/β-catenin/twist pathway. Cell Death and Disease, 10(3). 10.1038/s41419-019-1449-9

50. Méndez-Huergo, S. P., Blidner, A. G., & Rabinovich, G. A. (2017). Galectins: emerging regulatory checkpoints linking tumor immunity and angiogenesis. In Current Opinion in Immunology (Vol. 45, pp. 8–15). 10.1016/j.coi.2016.12.003

51. Banik, S. M., Pedram, K., Wisnovsky, S., Ahn, G., Riley, N. M., & Bertozzi, C. R. (2020). Lysosome-targeting chimaeras for degradation of extracellular proteins. Nature, 584(7820), 291–297. 10.1038/s41586-020-2545-9

52. Ebenezer, O., Comoglio, P., Wong, G. K. S., & Tuszynski, J. A. (2023). Development of Novel siRNA Therapeutics: A Review with a Focus on Inclisiran for the Treatment of Hypercholesterolemia. International Journal of Molecular Sciences, 24(4). 10.3390/ijms24044019

53. Mazella, J. (2001). Sortilin/neurotensin receptor-3: A new tool to investigate neurotensin signaling and cellular trafficking? In Cellular Signalling (Vol. 13, Issue 1, pp. 1–6). 10.1016/S0898-6568(00)00130-3

54. Rogers, H. T., Roberts, D. S., Larson, E. J., Melby, J. A., Rossler, K. J., Carr, A. V., Brown, K. A., & Ge, Y. (2023). Comprehensive Characterization of Endogenous Phospholamban Proteoforms Enabled by Photocleavable Surfactant and Top-down Proteomics. Analytical Chemistry. 10.1021/acs.analchem.3c01618

55. Larson, E. J., Pergande, M. R., Moss, M. E., Rossler, K. J., Wenger, R. K., Krichel, B., Josyer, H., Melby, J. A., Roberts, D. S., Pike, K., Shi, Z., Chan, H. J., Knight, B., Rogers, H. T., Brown, K. A., Ong, I. M., Jeong, K., Marty, M. T., McIlwain, S. J., & Ge, Y. (2023). MASH Native: a unified solution for native top-down proteomics data processing. Bioinformatics, 39(6). 10.1093/bioinformatics/btad359

56. Marty, M. T., Baldwin, A. J., Marklund, E. G., Hochberg, G. K. A., Benesch, J. L. P., & Robinson, C. V. (2015). Bayesian deconvolution of mass and ion mobility spectra: From binary interactions to polydisperse ensembles. Analytical Chemistry, 87(8), 4370–4376. 10.1021/acs.analchem.5b00140

57. Gilbert, L. A., Horlbeck, M. A., Adamson, B., Villalta, J. E., Chen, Y., Whitehead, E. H., Guimaraes, C., Panning, B., Ploegh, H. L., Bassik, M. C., Qi, L. S., Kampmann, M., & Weissman, J. S. (2014). Genome-Scale CRISPR-Mediated Control of Gene Repression and Activation. Cell, 159(3), 647–661. 10.1016/j.cell.2014.09.029

58. Hie, B. L., Shanker, V. R., Xu, D., Bruun, T. U. J., Weidenbacher, P. A., Tang, S., Wu, W., Pak, J. E., & Kim, P. S. (2023). Efficient evolution of human antibodies from general protein language models. Nature Biotechnology, 1–9. 10.1038/s41587-023-01763-2

