## Supporting Information for "A Genome-Wide CRISPR Screen Identifies Sortilin as the Receptor Responsible for Galectin-1 Lysosomal Trafficking"

Justin Donnelly, Roarke A. Kamber, Simon Wisnovsky, David S. Roberts, Egan L. Peltan, Michael C. Bassik, Carolyn R. Bertozzi

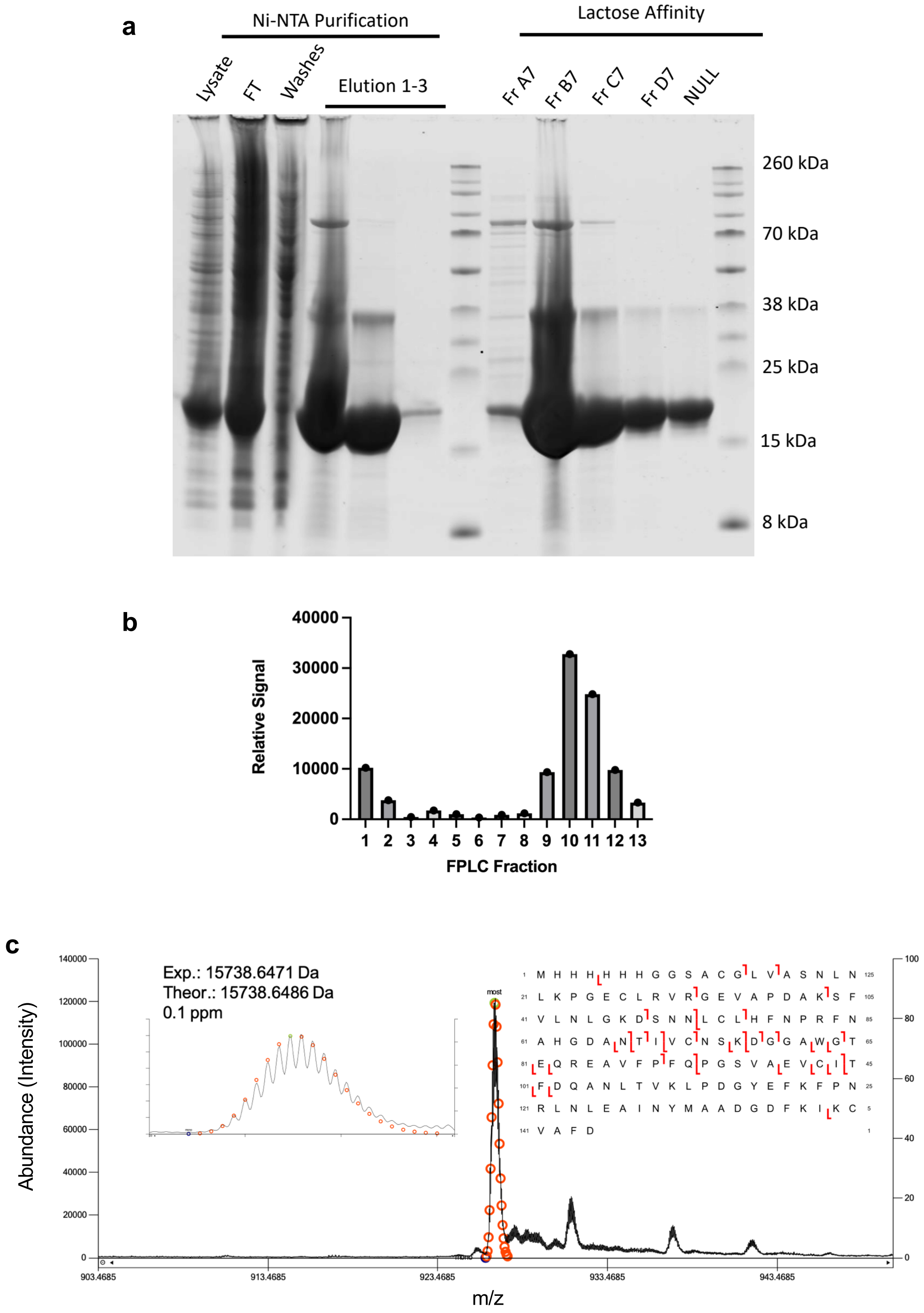


**Figure S1: Expression and Characterization of Gal1. (A)** SDS-PAGE of isolated fractions from E. coli lysates overexpressing 6xHis-tagged Gal1. Lysates were purified via Ni-NTA chromatography followed by lactose affinity FPLC. Representative fractions were analyzed via SDS-PAGE and total protein staining. **(B)** Quantification of Gal1 lactose affinity FPLC fractions determined via anti-6xHis Western blot. **(C)** Characterization of purified Gal1 by TDP-MS using MASH Native showing both the raw MS spectral alignment and the corresponding protein sequence table. Intact mass accuracy of the Gal1 most abundant molecular ion (z = 17+) was found to be within 0.1 ppm from the theoretical mass.

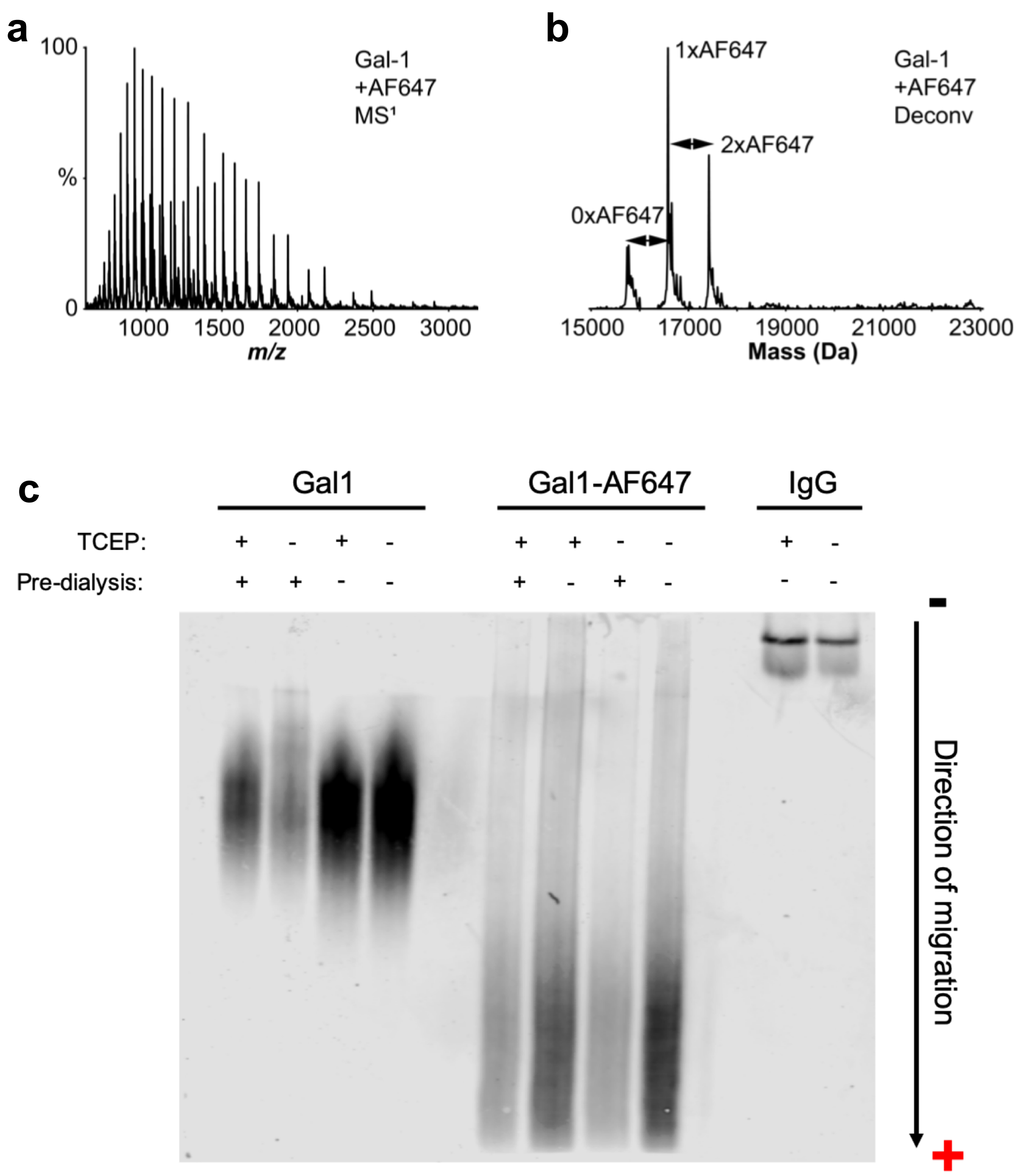


**Figure S2: Characterization of Gal1-AF647 by TDP-MS and Native PAGE. (A-B)** Raw (A) and deconvoluted (B) top-down mass spectra of Gal1-AF647. **(C)** Native PAGE of Gal1 and Gal1-AF647 stained with a total protein stain. Data indicate increase in native migration due to installation of AF647 dye consistent with the increased negative charge of this species.

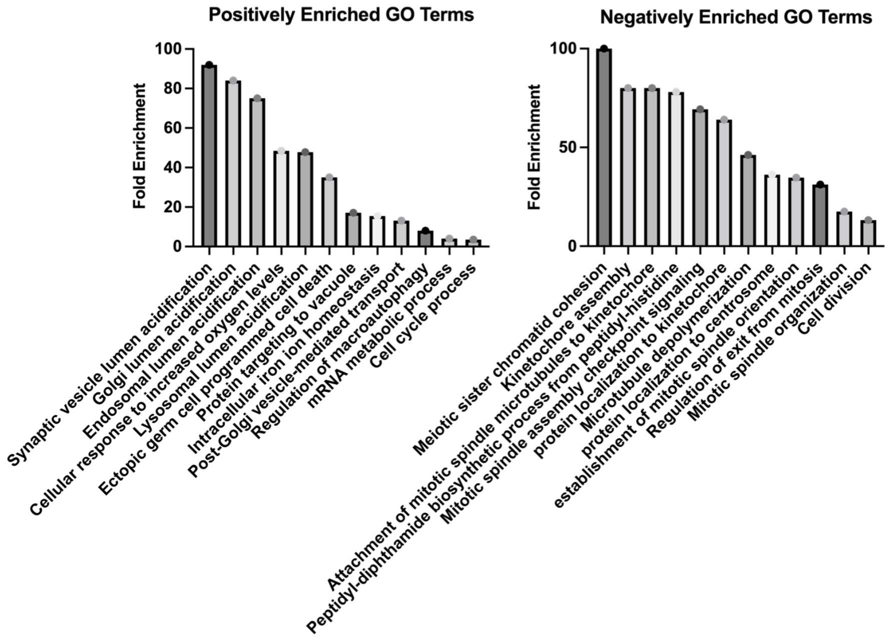


**Figure S3: GO Term Enrichment of Hits from Gal1 Internalization CRISPR Screen.** *Left:* Enriched GO terms for top 100 positively-enriched hits. *Right:* Enriched GO terms for top 100 negatively-enriched hits.

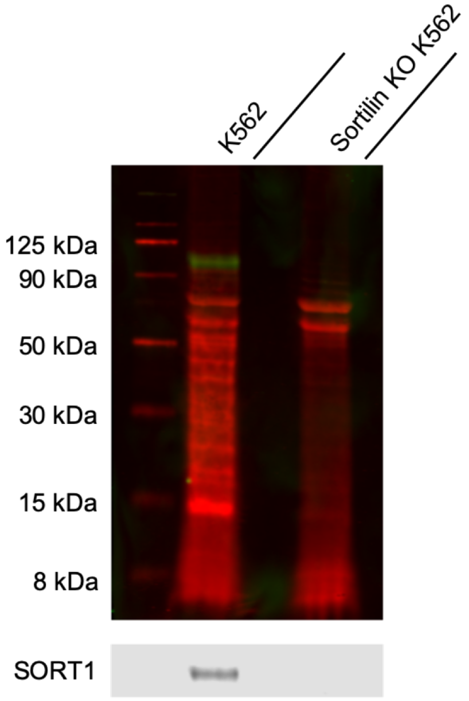


**Figure S4: Validation of *SORT1* KO in K562 cells.** *Top:* Western blot of WT and sortilin KO K562 cell lysates stained with total protein stain (red) and probed with anti-SORT1 (green). *Bottom:* Single channel anti-SORT1 stain.

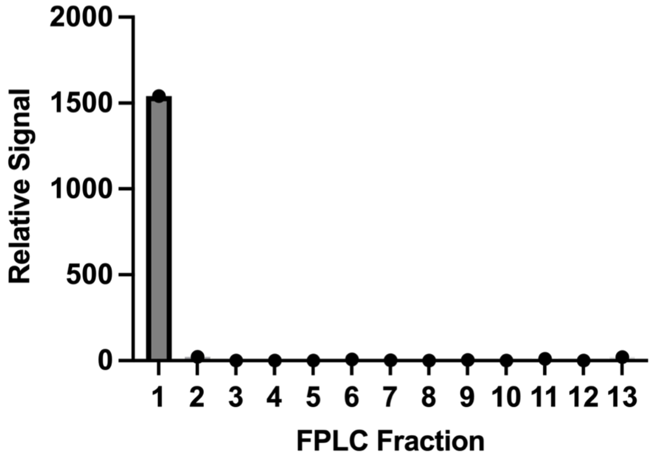


**Figure S5: Lactose affinity FPLC of Gal1(N46D).** Quantification of Gal1(N46D) lactose affinity FPLC fractions determined via anti-6xHis Western blot.

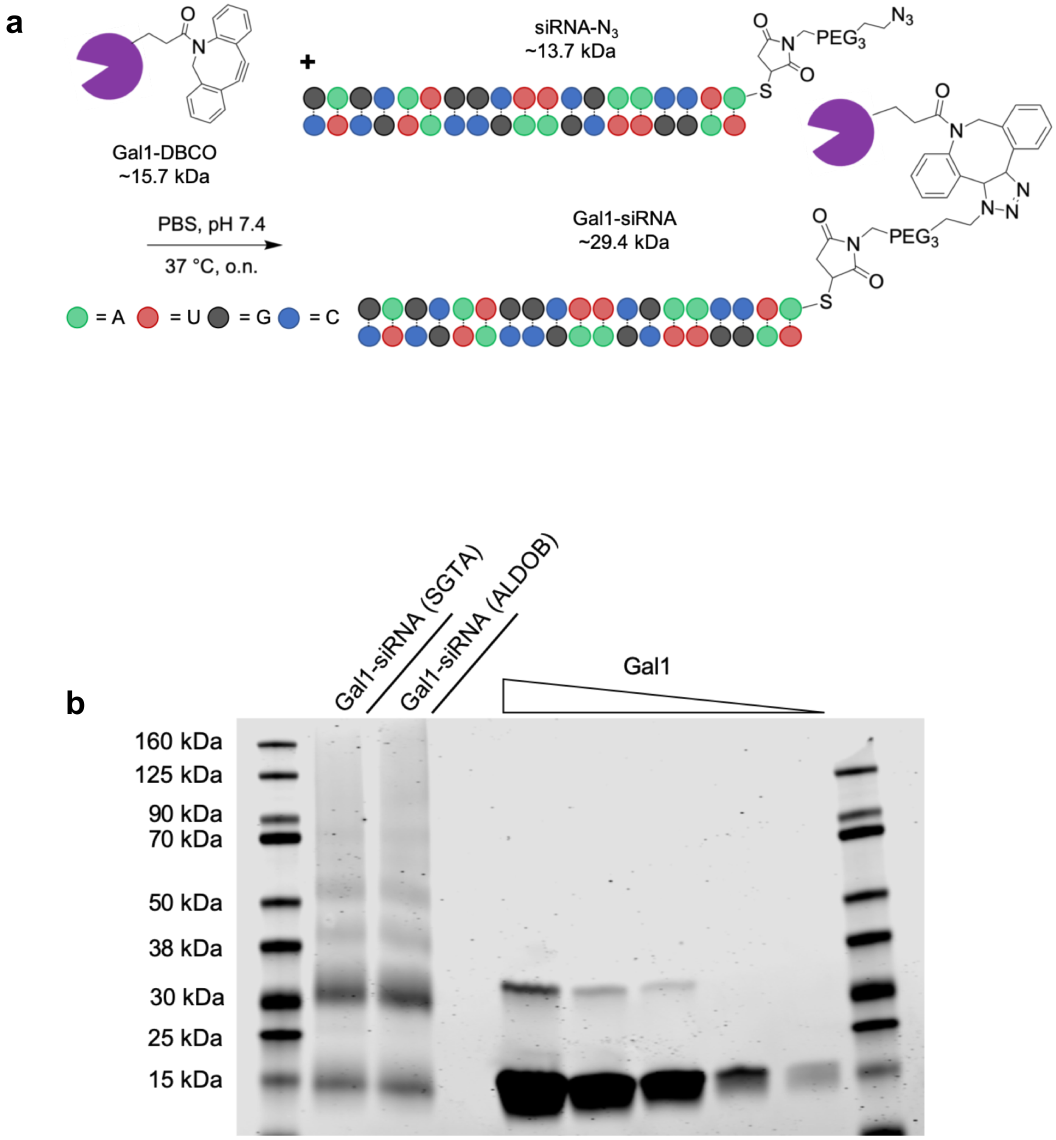


**Figure S6: Characterization of Gal1-siRNAs.** (A) Schematic of Gal1-siRNA preparation via click chemistry bioconjugation. DBCO-functionalized Gal1 ($PEG_4$ linker not shown) and azide-functionalized siRNA were combined to yield Gal1-siRNA conjugates. Estimated molecular weights also shown. (B) SDS-PAGE of two Gal1-siRNAs prepared via copper-free click chemistry. Gel was stained with a total protein stain. Five two-fold dilutions from 100 µg/mL unmodified Gal1 were also analyzed on the same gel for reference.

### Supplementary Experimental Methods

#### Protein Expression and Purification:

Plasmids obtained from Twist Bioscience (custom-ordered) were transfected into T7 Express cells (NEB, Cat. #C2566H) and grown on LB plates with appropriate selection marker. Single, sequence-confirmed clones were then inoculated into Terrific Broth (TB, ThermoFisher Cat. #22711022) with selection marker at starting OD 0.02-0.04. Clones were incubated at 37 °C until reaching OD 0.4-0.8 before induction of expression with 500 μM IPTG. Expression cultures were then transferred to 16 °C and incubated 48 hr. After pelleting at 4,000 xG for 5 min, cells were lysed via sonication on ice in lysis buffer (50 mM Tris-HCl + 300 mM NaCl + 5 mM $MgCl_2$ + 1 mM lactose + 10 mM imidazole + 1 mM TCEP + 0.1% Triton X-100 + 1X protease inhibitor cocktail (Thermo Scientific, Cat. # 78438) at pH 8.0). Lysates were clarified at ≥16,000 xG, 20 min, 4 °C. His-tagged proteins were subsequently purified via Ni-NTA agarose resin (Qiagen, Cat. #30210). After equilibration of resin with wash buffer (50 mM Tris-HCl + 300 mM NaCl + 5 mM $MgCl_2$ + 1 mM lactose + 10 mM imidazole + 1 mM TCEP + 0.05% Triton X-100 at pH 8.0) via three sequential washes (with centrifugation at 600 xG, 3 min), clarified lysates were added to resin and equilibrated for 15 min, 4 °C. Once equilibrated, resin was transferred to a purification column at 4 °C and washed with four column volumes of wash buffer. Sample was eluted with two column volumes of elution buffer (50 mM Tris-HCl + 300 mM NaCl + 5 mM $MgCl_2$ + 1 mM lactose + 250 mM imidazole + 1 mM TCEP + 0.025% Triton X-100 at pH 8.0). Fractions were analyzed via SDS-PAGE (see below for details). Where applicable, the lactose-binding fraction was isolated and analyzed using lactose-affinity FPLC. Protein solution was loaded onto a lactosyl-agarose column with a mobile phase of 50 mM Tris-HCl + 300 mM NaCl + 5 mM MgCl2 + 1 mM TCEP + 0.125% Triton X-100 at pH 8.0 and a gradient from 50 μM to 10 mM lactose. Protein quantification was accomplished via Pierce bicinchoninic acid (BCA) assay (ThermoFisher, Cat. #23227) according to manufacturer protocol.

#### Gel Electrophoresis and Western Blot:

Purified proteins, protein conjugates, and cell lysates were resolved on 4-12% XT Bis-Tris SDS-PAGE gels (BioRad, Cat. #3450123, 3450124) at 120-180 V in XT-MES buffer (BioRad, Cat. #1610789). Native PAGE was carried out on 8-16% TGX native gels (BioRad, Cat. #5671104) at 180 V, 4 °C in Tris-glycine buffer (Thermo Scientific, Cat. #LC2672). After electrophoresis, proteins were either stained with Bulldog Aquastain total protein (Bulldog Bio, Cat. #AS001000) for analysis or transferred to nitrocellulose membrane (BioRad, Cat #1620167). Where applicable, nitrocellulose membranes were stained with Revert total protein stain (LiCor, Cat. #926-11021) according to the manufacturer protocol and/or probed with primary antibody (1:1,000 in PBS-T) at 4 °C, overnight, followed by staining with corresponding fluorescent secondary antibody (1:10,000 in PBS-T) (purchased from LiCor) at r.t., 1 hr. Gels and membranes were analyzed using a LiCor Odyssey CLx imager.

**Chemicals:**

MMAF-DBCO was purchased from MedChem Express (Cat. #HY-133492) and MMAF was purchased from InVivo Chem (Cat. #V19682). All other click reagents were purchased from Click Chemistry Tools. AlexaFluor 647-NHS was purchased from Thermo Fisher Scientific (Cat. #A20006) and biotin-NHS was purchased from BroadPharm (Cat. #BP-22106). Recombinant human sortilin (carrier-free) was purchased from R&D Systems (Cat. #3154-ST-050). All other chemicals were purchased from various suppliers based on availability.

**Antibodies:**

The antibodies used in this work are identified in Table S1 below:

**Table S1: Antibodies**

| Target: | Supplier: | Catalogue Number: |
|---|---|---|
| **Gal1** | Abcam | Ab25138 |
| **Sortilin** | Sigma-Aldrich | HPA006889 |
| **Calnexin** | Thermo Scientific | PA5-34754 |
| **Calreticulin** | Thermo Scientific | PA3-900 |
| **GM130** | Thermo Scientific | PA1-077 |
| **LAMP1** | Thermo Scientific | PA1-654A |
| **EEA** | Thermo Scientific | PA1-063A |
| **IGF2R** | Thermo Scientific | PA3-850 |
